# A selective glucocorticoid receptor modulator attenuates lung inflammation and improves alveolarization in a neonatal rat model of bronchopulmonary dysplasia

**DOI:** 10.1101/2021.04.28.441723

**Authors:** Shoichi Ishikawa, Tohru Ogihara, Shigeo Yamaoka, Jun Shinohara, Shigeru Kawabata, Yoshinobu Hirose, Akira Ashida

## Abstract

**Background:** Bronchopulmonary dysplasia (BPD) is a major problem for extremely preterm infants. Glucocorticoids effectively treat BPD; however, they have many side effects. Compound A (Cpd A) is a nonsteroidal Selective Glucocorticoid Receptor Modulator (SEGRM) that acts as a glucocorticoid receptor ligand without inducing the expression of glucocorticoid-response element-driven genes. Cpd A reportedly has anti-inflammatory properties with fewer side effects than glucocorticoids.

**Methods:** Using a bleomycin (Bleo)-induced BPD model, we evaluated the therapeutic effects of Cpd A. 0-day-old Sprague-Dawley rats were administered Bleo for 10 days and treated with dexamethasone (Dex) or Cpd A from day 0 to 13. We evaluated lung pathology by histology and the mRNA levels of interleukin (IL)-1β, transforming growth factor (TGF)-β1 and chemokines, CXCL1 and CCL2.

**Results:** Bleo-treated mice had lungs with impaired alveolarization. Dex and Cpd A treatments improved the alveolar structure, attenuating the lung injury. Bleo-exposed lungs had increased inflammatory cells recruitment and inflammatory mediator mRNA levels. Cpd A treatment reduced inflammatory cells infiltration and CXCL1, CCL2 and TGF-β1 expression.

**Conclusion:** Cpd A improved lung inflammation and alveolar maturation arrest, and restored histological and biochemical changes in a model of BPD. SEGRMs, including Cpd A, are promising candidates for the therapy of BPD.

**Impact Statement:**

○ What is the key message of your article? Compound A decreased lung inflammation and improved lung morphometric changes in Bleomycin-exposed lungs.
○ What does it add to the existing literature? Compound A has anti-inflammatory effects in an experimental model of BPD.
○ What is the impact? SEGRMs, including Cpd A, may be promising candidates for the therapy of BPD.

## INTRODUCTION

Advances in perinatal and neonatal care have increased the survival rate of extremely preterm infants (1). However, bronchopulmonary dysplasia (BPD) remains a major cause of morbidity for extremely preterm infants. One report suggests that 40 % of infants born at ≤ 28 weeks’ gestation experienced some degree of BPD (2). BPD is a multifactorial disease, and inflammation plays a key role in its development (3). BPD is characterized by inflammation and alveolar maturation arrest during the early stage and by wall thickness and fibrosis during the late stage of disease (4).

Glucocorticoids (GCs) have anti-inflammatory properties and show effectiveness for BPD (5). After GC receptors (GRs) bind to GCs, they regulate gene expression with two processes: transactivation and transrepression (6). In transactivation, a GR-homodimer binds to glucocorticoid-response elements (GREs) and induces the expression of anti-inflammatory genes (6) (Supplemental figure. 1). Conversely, during transrepression, GRs interfere with other transcription factors associating with inflammation, such as nuclear factor (NF)-κB or activator protein (AP)-1, and inhibit the expression of pro-inflammatory gene (6) (Supplemental figure. 1). This process does not require GR dimerization (6). However, GCs are known to have many side effects such as immunosuppression and diabetes. GCs can also cause a delay in somatic growth and impair neurodevelopment, which is particularly problematic in infants (7). Selective Glucocorticoid Receptor Modulators (SEGRMs) are nonsteroidal compounds. SEGRMs can act as a ligand for GRs and have anti-inflammatory effects; however, they do not induce GR dimerization (8, 9). SEGRMs are reported to have anti-inflammatory properties through transrepression (8, 9). Some side effects such as hyperglycemia and muscle wasting are due to byproducts during transactivation (8), and SEGRMs are expected to have fewer side effects than GCs. Several studies have indicated that SEGRMs do have fewer side effects than GCs (6, 8, 9).

Compound A (Cpd A), 2-(4-acetoxyphenyl)-2-chloro-N-methylethylammonium chloride, is a SEGRM. Cpd A has anti-inflammatory properties using only the transrepression pathway and is expected to have fewer side effects than GCs. There are several reports showing the efficacy of Cpd A in various inflammatory models, such as collagen-induced arthritis (10, 11), muscular dystrophy (12), or a Th2-driven asthma model (13). They also showed that Cpd A did not have the severe adverse effects reported for GCs, including hyperglycemia (9), osteoporosis (14), or muscle degradation (12).

In this study, we investigated the effects of Cpd A in a rat model of BPD. We used a bleomycin-induced neonatal rat model, which is well established as a useful model of the alveolar maturation arrest seen in BPD (4). Using this BPD model, we evaluated the effect of Cpd A on lung inflammation and alveolar maturation arrest as well as histological and biochemical alterations. We hypothesized that Cpd A would attenuate lung inflammation and improve lung morphometric changes in an experimental model of BPD.

## METHODS

All procedures were approved by the research animal committee of Osaka Medical Collage (authorized number 29087, 30091 and 2019-092).

### Subjects

Pregnant Jcl/SD rats were purchased from CREA Japan, Inc. (Tokyo, Japan). Pups were delivered naturally (on day 21) and were nursed by the dam. Animals were fed ad libitum and exposed to 12:12-h light-dark cycles throughout the study period. Bleomycin sulfate (Bleo) was purchased from LKT Laboratories, Inc. (St. Paul, MN). Dexamethasone (Dex) was purchased from Fujifilm Wako Pure Chemical Corporation Inc. (Osaka, Japan). Compound A (Cpd A) was purchased from Santa Cruz Biotechnology, Inc. (Santa Cruz, CA). Anti-cluster of differentiation (CD) 68 antibody (catalog no. ab31630) was purchased from Abcam, Inc. (Cambridge, UK). Anti-Ly6G antibody (catalog no. HM1039) was purchased from Hycult Biotech Inc. (Wayne, PA).

### Protocol

Zero-day-old rats from timed-pregnant dams were divided into four groups; 1) Control group: Normal saline-exposed and vehicle (distilled water)-treated, 2) Bleo group: Bleo-exposed (1 mg/kg) and vehicle-treated, 3) Dex group: Bleo-exposed and Dex-treated (0.1 mg/kg), and 4) Cpd A group: Bleo-exposed and Cpd A-treated (1 mg/kg). All reagents were administered by intraperitoneal injection. Bleo or normal saline were injected from day 0 to day 10. Vehicle, Dex, or Cpd A were injected until day 13. On day 10 or day 14 of life, pups were euthanized with an intraperitoneal injection of pentobarbital sodium (100 mg/kg) (Supplemental figure. 2).

### Fixation of lung tissue

After the rats were euthanized, their left lungs were removed and snapped frozen in liquid nitrogen, and stored separately at −80°C. They were used for subsequent RNA analysis. The right lung was air-inflated and perfusion-fixed at constant pressure, removed along with the heart, and fixed in 4 % paraformaldehyde. They were embedded in paraffin and used for subsequent histological analysis. Paraffin embedded lung tissues were cut into 4 μm sections for subsequent histological analyses of lung injury and immunohistochemical analysis.

### Lung morphometry and alveolar development

The right lungs obtained on day 14 of life were used. The sections were stained with hematoxylin and eosin and the radial alveolar count (RAC) and mean linear intercept length (MLI) were measured. Each section was captured on a Nikon Eclipse 80i microscope, using the 10× objective and captured by DS-Ri1 digital camera (Nikon, Tokyo, Japan). At least 10 random lung fields were photographed from each animal. Measurements were carried out on three noncontiguous right lung sections per animal by a single observer blinded to the group identity.

The RAC was determined as previously described (4, 15). Respiratory bronchioles were identified as bronchioles lined by epithelia in one part of the wall. From the center of the respiratory bronchiole, a perpendicular line was drawn to the edge of the acinus (as defined by a connective tissue septum or the pleura), and the number of septa intersected by this line was counted. The MLI was measured as previously reported (4, 16). Large airways and vessels were avoided. Grids of horizontal and vertical lines were superimposed on an image and the number of times the lines intersected with the tissue was counted. The total length of the grid lines was then divided by the number of intersections to provide the MLI in μm.

### Immunohistochemical analysis

Lung tissue was immunostained for CD68 to identify macrophages and for Ly6G to identify neutrophils. The right lungs at day 14 of life were used. Paraffin-embedded slides from formalin-fixed tissue were deparaffinized in xylene. The sections were rehydrated by serial immersions in 100 % ethanol, 90 % ethanol, 70 % ethanol, and water. Antigen retrieval was conducted with a microwave oven and then washed with phosphate buffered saline (PBS). Endogenous peroxidase activity was reduced by immersion in 3 % hydrogen peroxide. After rinsing, sections were covered with 3 % goat serum for 30 min and incubated with anti-CD68 antibody diluted in PBS (1:800) and with anti-Ly6G antibody diluted in PBS (1:400) overnight. After incubation, the sections were rinsed with PBS and incubated with biotin-labeled secondary antibody diluted 1:300 in PBS for 2 h. After incubation with the secondary antibody, the sections were rinsed with PBS, incubated in avidin-biotin complex staining kit (Vector Laboratories, Burlingame, CA) for 30 min at room temperature, rinsed in PBS, and developed with diaminobenzidine and hydrogen peroxide. Slides were lightly counterstained with hematoxylin. The slides were then dehydrated by sequential immersion in 70 % ethanol, 90 % ethanol, 100 % ethanol, and xylene before applying coverslips. Analysis of tissue macrophage (CD68-positive) and neutorophil (Ly6G-positive) numbers was conducted from 10 random, non-overlapping, high-power fields captured from each section. Measurements were conducted on three noncontiguous right lung sections per animal by two observers blinded to the group identity.

### Real-time PCR

The left lung tissue was used to analyze gene expression at day 10 (to evaluate lung inflammation soon after Bleo exposure) and day 14. Total RNA was extracted with ISOGEN (Nippon Gene, Tokyo, Japan), according to the manufacturer’s protocol. The RNA purity was verified by A260/A280 ratio. Total RNA was reverse transcribed into cDNA using an Omniscript Reverse Transcription Kit (Qiagen, Venlo, Netherlands). The mRNA levels of cytokines and chemokines were measured with reverse transcription–polymerase chain reaction (RT-PCR) with TaqMan using a StepOnePlus real time PCR system (Thermo Fisher Scientific, Waltham, MA). We analyzed the mRNA expression level of interleukin-1β (IL-1β; catalog no. Rn99999009_m1), C-X-C motif chemokine ligand 1 (CXCL1; catalog no. Rn00578225_m1), C-C motif chemokine 2 (CCL2; catalog no. Rn00580555_m1), and transforming growth factor-β1 (TGF-β1; catalog no. Rn00572010_m1). Glyceraldehyde-3-phosphate dehydrogenase (GAPDH; catalog no. Rn01775763_g1) was used as a standardizing control. These TaqMan primers (TaqMan Gene Expression Assays) were purchased from Thermo Fisher Scientific Inc. (Waltham, MA). We chose IL-1β as a major inflammatory cytokine and CCL2 as a potent chemoattractant for inflammatory cells. We also chose CXCL1 as a neutrophil attractant, and TGF-β1 as one of the major growth factors associated with fibrosis. The expression level of each mRNA was calculated using the 2^− ΔΔCT^ method.

### Statistics

All statistical analyses were carried out using JMP Pro Version 14.0 (SAS Institute Inc., Cary, NC). Data were expressed as mean ± SE and analyzed by Wilcoxon signed-rank test. A significant value was considered to be *p*<0.05.

## RESULTS

### Body weight

Of the 56 animals studied, three animals died during the study: one in the Bleo group, one in the Dex group and one in the Cpd A group.

Pups treated with Dex had significantly lower body weights on day 14 compared with Control and Bleo groups (33.4±2.2 g vs 33.9±1.7 g vs 24.7 ±2.0 g, Control vs Bleo vs Dex; *p*<0.05, Fig. 1). The body weights in the remaining three groups were not statistically different to one another.

**Figure 1.**
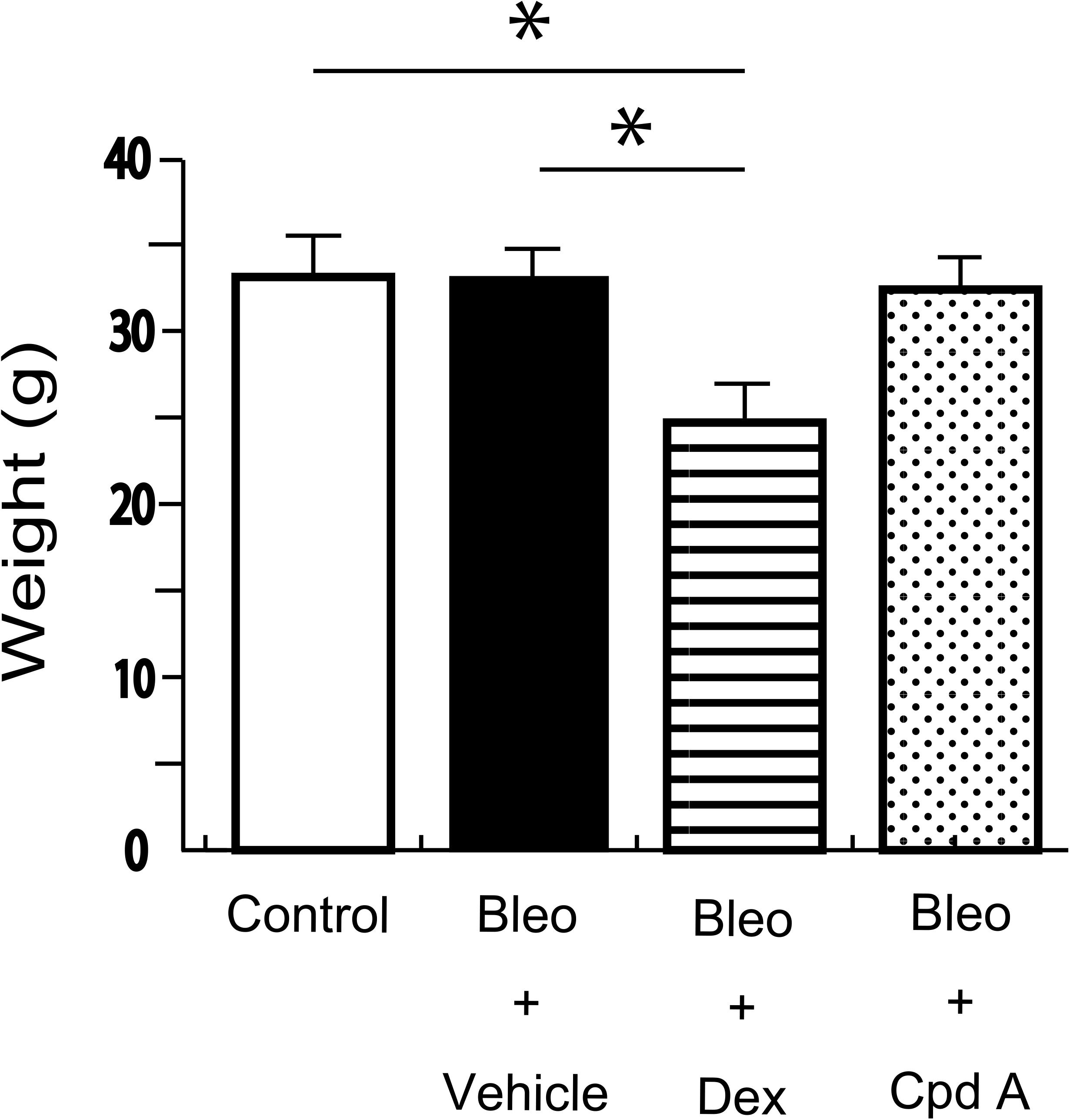
Body weight at day 14 of life. Rats in Dex group at 14 days of age had significantly lower body weight compared with Control and Bleo groups (33.4±2.2 g vs 33.9±1.7 g vs 24.7 ±2.0 g, Control vs Bleo vs Dex; *p*<0.05; N=5-8/group). The body weights in the remaining 3 groups were not statistically different to one another. White bar indicates Control group, black bar indicates Bleo group, horizontal striped bar indicates Dex group and polka dot bar indicates Cpd A group. One asterisk signifies a significance level of p<0.05, two asterisks signify a significance level of p<0.01.

### Morphometric analysis

The lungs from the Bleo group displayed impaired alveolarization as demonstrated by decreased septation, distal airspace enlargement, and a reduction in complexity compared with the Control group (Fig. 2A, B). Treatment with Dex or Cpd A improved the alveolar structure and attenuated the lung injury (Fig. 2C, D). These differences were assessed by MLI and RAC. MLI was significantly increased in rats exposed to Bleo compared with control animals (69.5±3.6 μm vs 120.7±3.0 μm, Control vs Bleo; *p*<0.01, Fig. 2E). The Dex and Cpd A groups had significantly lower MLI compared with the Bleo group (120.7±3.0 μm vs 92.9±3.0 μm vs 95.7±2.8 μm, Bleo vs Dex vs Cpd A; *p*<0.01, Fig. 2E); however, the MLI values remained higher than controls. RACs were significantly reduced in rats exposed to Bleo compared with control animals (13.3±0.2 vs 7.9±0.2, Control vs Bleo; *p*<0.01, Fig. 2F). The Dex and Cpd A groups had significantly higher RAC compared with the Bleo group (7.9±0.2 vs 11.6±0.2 vs 12.1±0.2, Bleo vs Dex vs Cpd A; *p*<0.01, Fig. 2F); however, the RAC values remained lower than controls.

**Figure 2.**
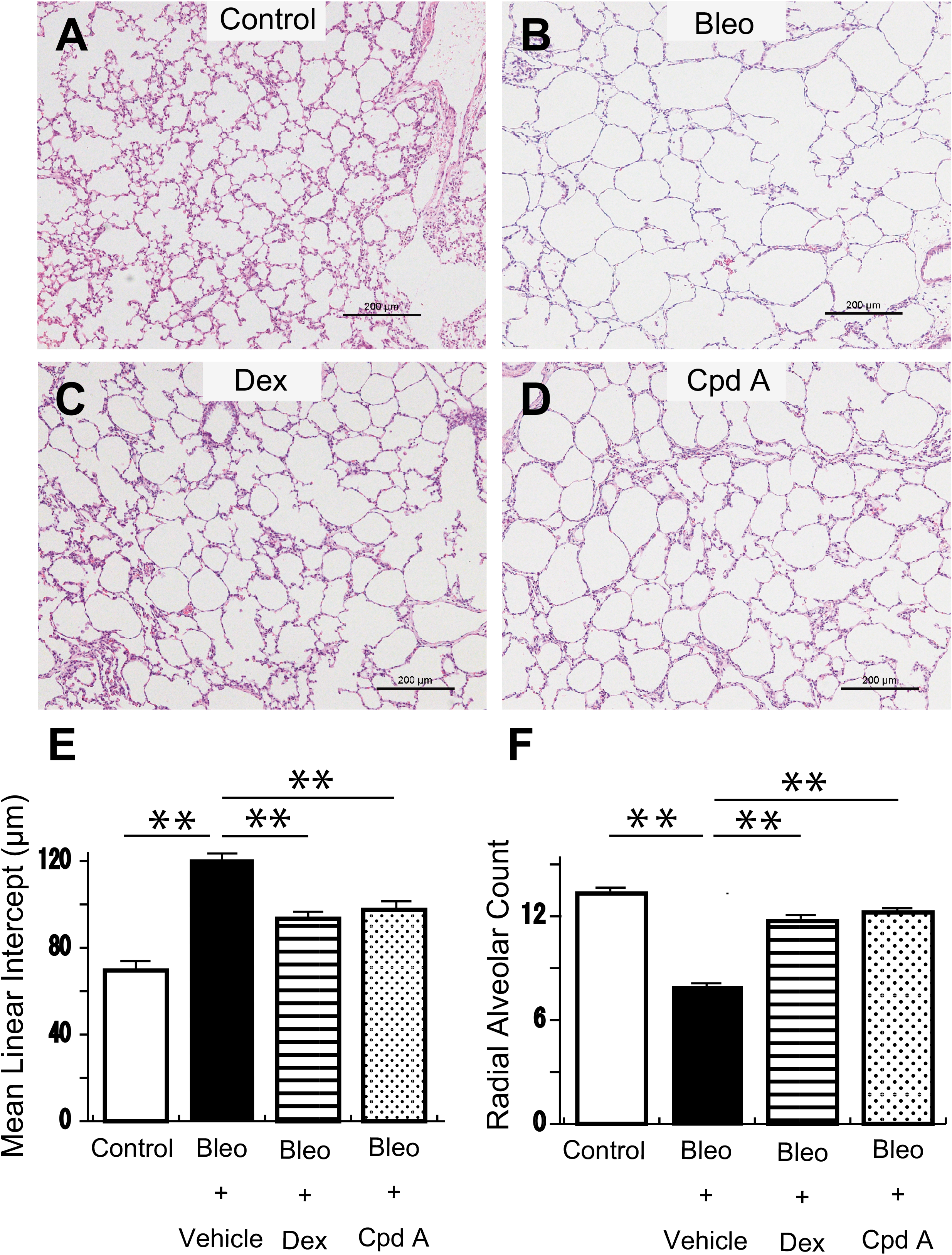
Mean linear intercept (MLI) and Radial alveolar counts (RACs) in newborn rats exposed to Bleo for 13 days. Lung histology from **A** Control group, **B** Bleo group, **C** Dex group, and **D** Cpd A group at day 14 of life, stained with hematoxylin and eosin. Magnification × 100. Scale bar is 200μm. **E** Compared with Control group, MLI increased in Bleo-exposed rats (69.5±3.6 μm vs 120.7±3.0 μm, Control vs Bleo; *p*<0.01; N=5-7/group). MLI of both Dex-treated and Cpd A-treated rats decreased compared with Bleo group (120.7±3.0 μm vs 92.9±3.0 μm vs 95.7±2.8 μm, Bleo vs Dex vs Cpd A; *p*<0.01; N=7-8/group). **F** Compared with control, RAC decreased in Bleo-exposed rats (13.3±0.2 vs 7.9±0.2, Control vs Bleo; *p*<0.01; N=5-7/group). RAC of both Dex-treated and Cpd A-treated rats increased compared with Bleo group (7.9±0.2 vs 11.6±0.2 vs 12.1±0.2, Bleo vs Dex vs Cpd A; *p*<0.01; N=7-8/group). White bar indicates Control group, black bar indicates Bleo group, horizontal striped bar indicates Dex group and polka dot bar indicates Cpd A group. One asterisk signifies a significance level of p<0.05, two asterisks signify a significance level of p<0.01.

Lungs from the rats treated with Dex or Cpd A alone following normal saline-exposure were also evaluated. Neither group showed significant morphometrical changes compared with the control group (data not shown).

### Immunohistochemical analysis

The Bleo-exposed newborn rats showed increased recruitment of macrophages to their lungs as determined by immunohistochemical analysis (34.7±3.3 cells/10HPF vs 46.3±1.7 cells/10HPF, Control vs Bleo; *p*<0.01, Fig. 3A, B, and I). Furthermore, the number of macrophages was significantly reduced following Cpd A treatment compared with Bleo group (46.3±1.7 cells/10HPF vs 40.2±1.8 cells/10HPF, Bleo vs Cpd A; *p*<0.05, Fig. 3B, D, and I). There was no difference between Bleo group and Dex group (46.3±1.7 cells/10HPF vs 45.7±2.0 cells/10HPF, Bleo vs Dex; *p*=0.84, Fig. 3B, C, and I).

**Figure 3.**
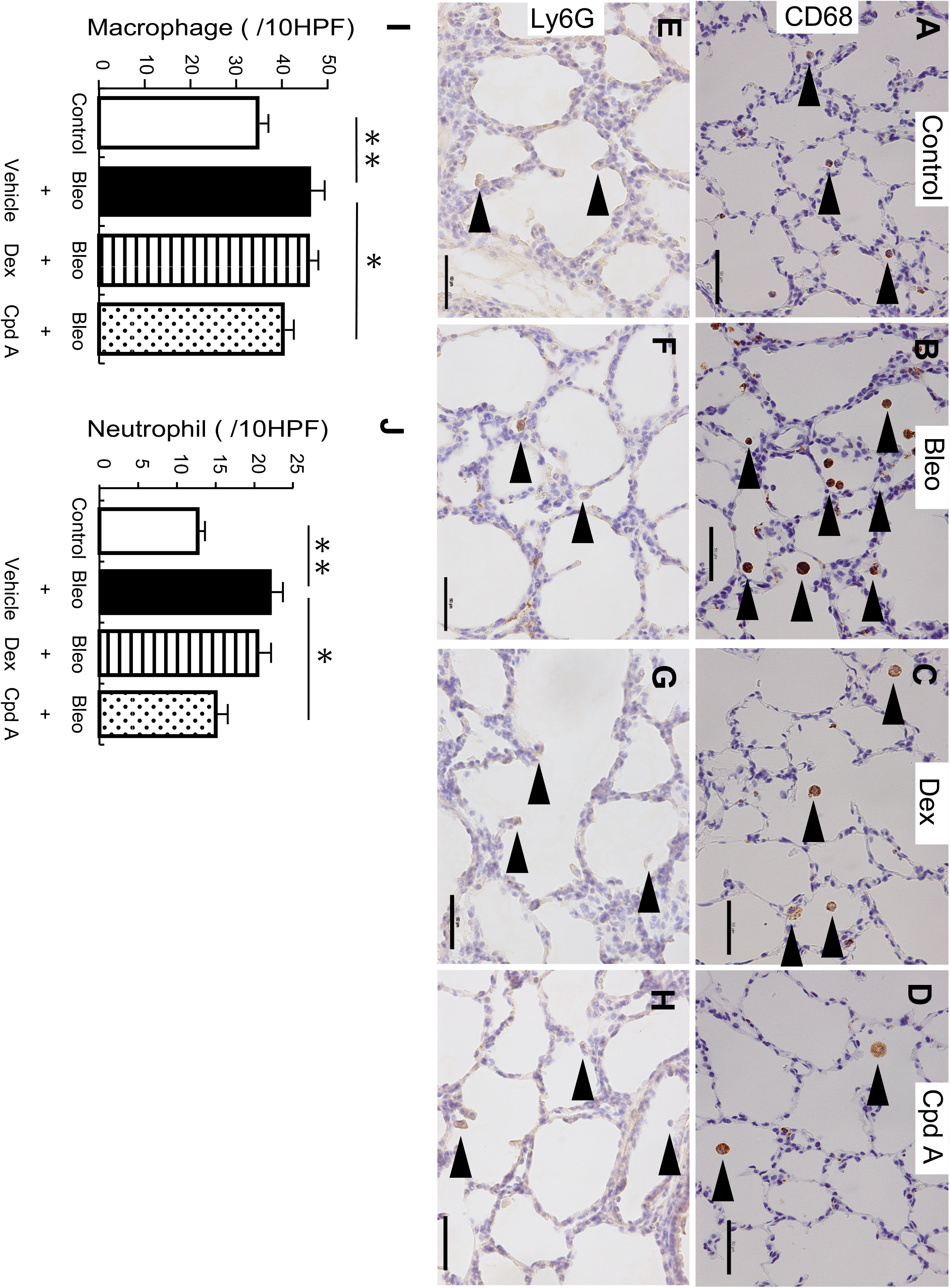
Cpd A decreased the number of macrophages and neutrophils infiltrating lungs in experimental BPD. **A-D** anti-CD68 immunostaining, **A** Control group, **B** Bleo group, **C** Dex group, and **D** Cpd A group. **E-H** anti-Ly6G immunostaining, **E** Control group, **F** Bleo group, **G** Dex group, and **H** Cpd A group. Magnification × 400. Scale bar is 50μm. Brown stained cells are CD68 or Ly6G positive (arrow). **I** The recruitment of macrophages increased in the Bleo-exposed rats compared with Control group (34.7±3.3 cells/10HPF vs 46.3±1.7 cells/10HPF, Control vs Bleo; *p*<0.01; N=5-9/group). The number of macrophages was significantly reduced following Cpd A treatment (46.3±1.7 cells/10HPF vs 40.2±1.8 cells/10HPF, Bleo vs Cpd A; *p*<0.05; N=8-9/group). There was no difference between Bleo group and Dex group (46.3±1.7 cells/10HPF vs 45.7±2.0 cells/10HPF, Bleo vs Dex; *p*=0.84; N=7-9/group). **J** The recruitment of neutrophils in the Bleo-exposed lungs also increased (12.7±1.4 cells/10HPF vs 22.1±1.4 cells/10HPF, Control vs Bleo; *p*<0.01; N=8-9/group). The number of neutrophils was significantly reduced following Cpd A treatment compared with Bleo (22.1±1.4 cells/10HPF vs 15.0±1.4 cells/10HPF, Bleo vs Cpd A; *p*<0.05; N=8-9/group). There was no difference between Bleo and Dex (22.1±1.4 cells/10HPF vs 20.4±1.5 cells/10HPF, Bleo vs Dex; *p*=0.36; N=8-9). White bar indicates Control group, black bar indicates Bleo group, horizontal striped bar indicates Dex group and polka dot bar indicates Cpd A group. One asterisk signifies a significance level of p<0.05, two asterisks signify a significance level of p<0.01.

Neutrophil recruitment in Bleo-exposed lungs also increased (12.7±1.4 cells/10HPF vs 22.1±1.4 cells/10HPF, Control vs Bleo; *p*<0.01, Fig. 3E, F, and J). Furthermore, the number of neutrophils was significantly reduced following Cpd A treatment compared with the Bleo group (22.1±1.4 cells/10HPF vs 15.0±1.4 cells/10HPF, Bleo vs Cpd A; *p*<0.05, Fig. 3F, H, and J). There was no significant difference between Bleo and Dex groups (22.1±1.4 cells/10HPF vs 20.4±1.5 cells/10HPF, Bleo vs Dex; *p*=0.36, Fig. 3F, G, and J).

### mRNA expression of cytokines and chemokines

In the Bleo-exposed lungs on day 10 of life, all inflammatory mediators demonstrated a significant increase in gene expression compared with Control group (*p*<0.05; Control vs Bleo, Fig.4A, B, and C). In particular, CXCL1 and CCL2 showed a 2-fold or greater increase.

Cpd A led to a significant decrease of CXCL1 and CCL2 gene expression compared with Bleo group (*p*<0.05; Bleo vs Cpd A, Fig. 4B, C). Cpd A treatment also decreased IL-1β expression; however, this change was not statistically significant (*p*=0.06; Bleo vs Cpd A, Fig. 4A).

**Figure 4.**
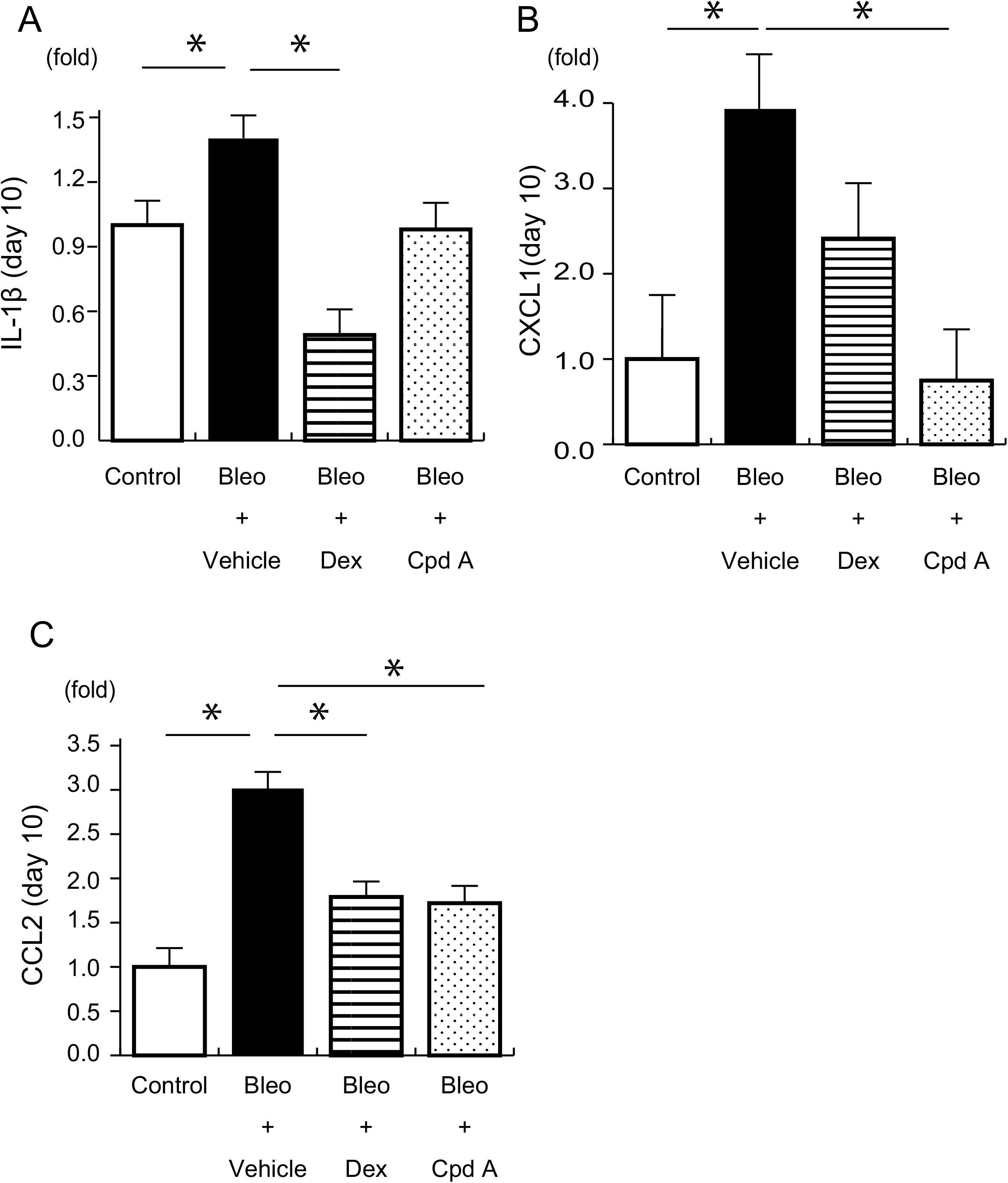
The mRNA expression of inflammatory mediators at day 10 of life. **A** IL-1β, **B** CXCL1, and **C** CCL2. All mediators increased in Bleo-exposed rats compared with Control (*p*<0.05; Control vs Bleo; N=3-4/group). Compared with Bleo-exposed rats, the expression of CXCL1 and CCL2 decreased in Cpd A-treatment rats (*p*<0.05; Bleo vs Cpd A; N=4-5/group). The expression of IL-1β also decreased in Cpd A group, but this change was not statistically significant (*p*=0.06; Bleo vs Cpd A; N=4/group). Compared with Bleo group, Dex led to a significant decrease of IL-1 β and CCL2 in gene expression (*p*<0.05; Bleo vs Dex; N=4-5/group). There was no difference in CXCL1 gene expression between Bleo group and Dex group (*p*=0.22; Bleo vs Dex; N=4/group). White bar indicates Control group, black bar indicates Bleo group, horizontal striped bar indicates Dex group and polka dot bar indicates Cpd A group. One asterisk signifies a significance level of p<0.05, two asterisks signify a significance level of p<0.01.

Compared with Bleo group, Dex led to a significant decrease in IL-1β and CCL2 gene expression (*p*<0.05; Bleo vs Dex, Fig. 4A, C). There was no difference in CXCL1 gene expression between Bleo and Dex group (*p*=0.22; Bleo vs Dex, Fig. 4B).

On day 14 of life, the expression of IL-1β was decreased compared with Control group in Bleo-exposed lungs (*p*<0.05; Control vs Bleo, Fig. 5A). The expression of the other inflammatory mediators showed no difference between the Bleo-exposed rats and Control group (Control vs Bleo, Fig. 5B, C). Furthermore, Dex did not influence the expression of these inflammatory mediators (Bleo vs Dex, Fig. 5A, B, and C), while Cpd A treatment decreased IL-1β compared with Control and Dex groups (*p*<0.05; Control vs Dex vs Cpd A, Fig. 5A).

**Figure 5.**
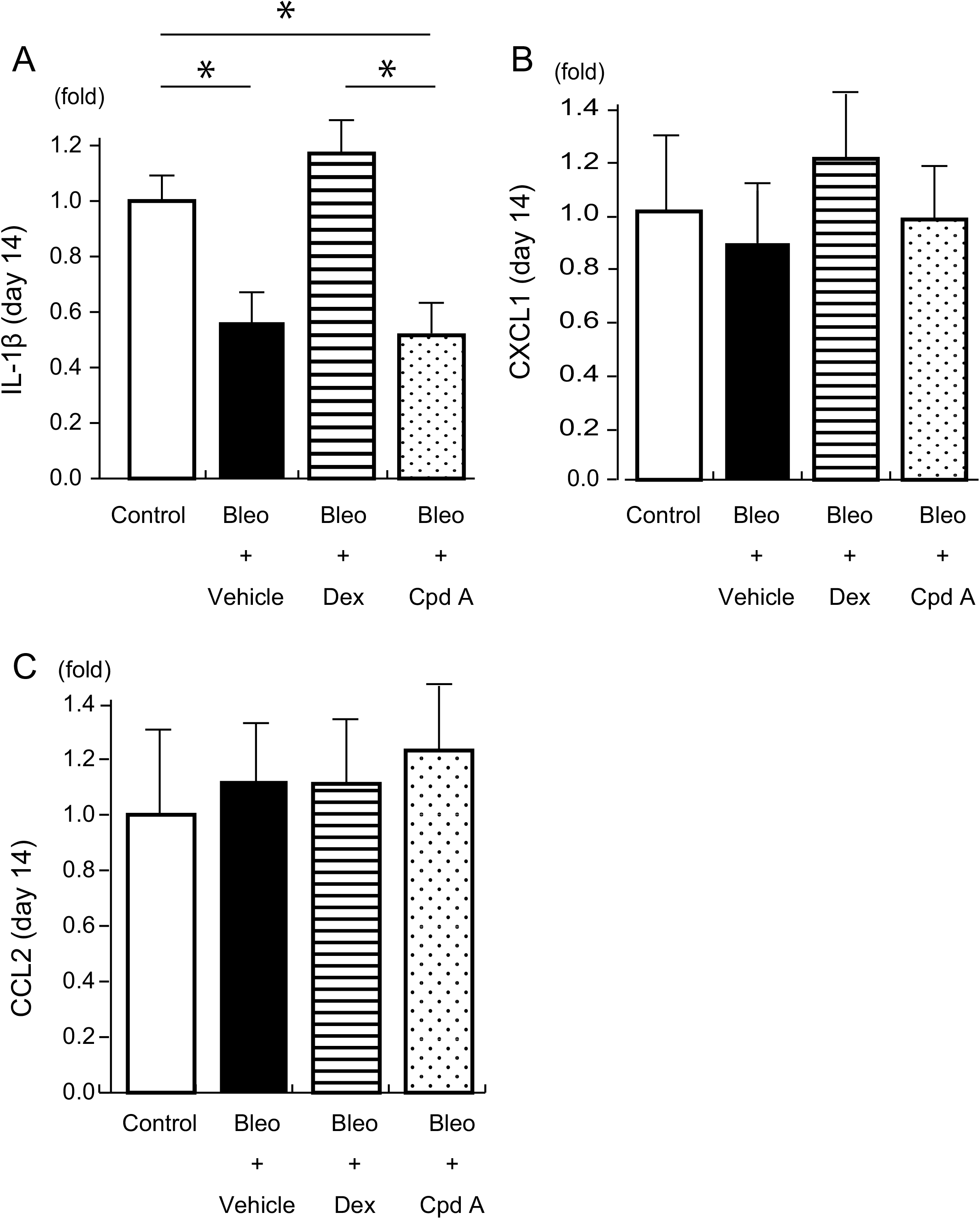
The mRNA expression of inflammatory mediators at day 14 of life. **A** IL-1β, **B** CXCL1, and **C** CCL2. **A** The expression of IL-1β decreased compare with Control group in Bleo-exposed lungs *(p*<0.05; Control vs Bleo; N=4-8/group). **B C** The expression of the other inflammatory mediators showed no difference between the Bleo-exposed rats and Control group (Control vs Bleo; N= 4-8/group). Dex did not change the gene expression of these inflammatory mediators (Bleo vs Dex; N=7-8/group), while Cpd A treatment decreased IL-1β compared with Control and Dex group (*p*<0.05; Control vs Dex vs Cpd A, N=4-8/group). White bar indicates Control group, black bar indicates Bleo group, horizontal striped bar indicates Dex group and polka dot bar indicates Cpd A group. One asterisk signifies a significance level of *p*<0.05, two asterisks signify a significance level of p<0.01.

The mRNA expression of TGF-β1 was also analyzed on day 10 and 14 of life (Fig. 6A, B). On day 10, TGF-β1 was lower in Bleo-exposed rats lungs compared with the controls (*p*<0.05; Control vs Bleo, Fig. 6A). Conversely, on day 14, TGF-β1 expression was increased significantly in the Bleo group (*p<* 0.05; Control vs Bleo, Fig. 6B). The increase in TGF-β1 expression in Bleo-exposed lungs was significantly suppressed by both the Dex and Cpd A groups (*p*<0.05; Bleo vs Dex vs Cpd A, Fig. 6B).

**Figure 6.**
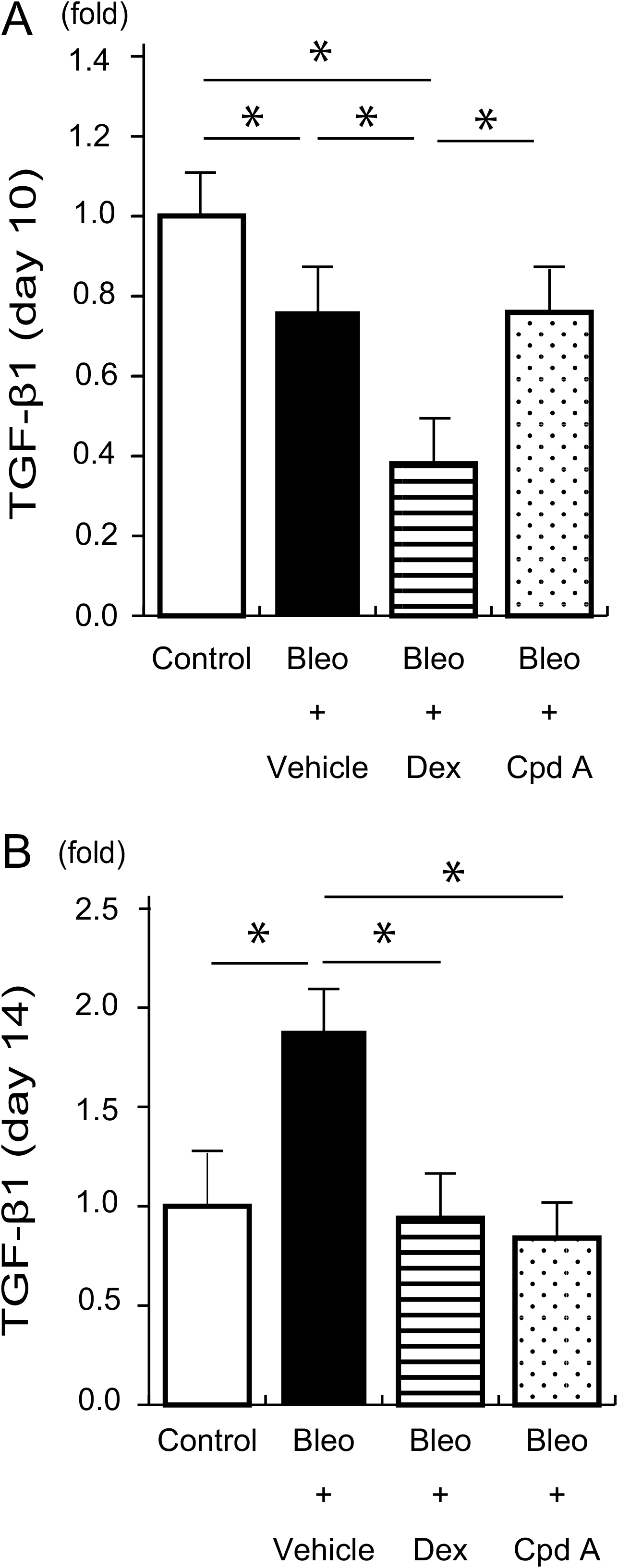
The mRNA expression of TGF-β1 at day 10 and 14. **A** The level of TGF-β1 in the Bleo-exposed rats at 10 days of age decreased relatively compared with control(*p*<0.05; Control vs Bleo; N= 3-4/group). **B** At day 14, TGF-β1 expression was increased significantly in the Bleo group. The increase in TGF-β1 expression in Bleo-exposed lungs was suppressed by Dex and Cpd A (*p*<0.05; Bleo vs Dex vs Cpd A; N=5-7/group). White bar indicates Control group, black bar indicates Bleo group, horizontal striped bar indicates DEX group and polka dot bar indicates Cpd A group. One asterisk signifies a significance level of p<0.05, two asterisks signify a significance level of p<0.01.

## DISCUSSION

In the present study, treatment with Dex or Cpd A improved morphometrical changes in the Bleo-exposed lungs. The exposure of neonatal rats to bleomycin disrupts the normal lung structure, as evidenced by decreased septation, distal airspace enlargement, and reduction in complexity. In general, BPD animal models do not capture the entire disease, but model a pathological event (17). Our animal model represented alveolar growth arrest and is recognized as an alveolar simplification model as seen in human BPD. Treatment with Dex or Cpd A improved these morphometrical changes.

The mRNA expression of inflammatory mediators significantly increased in Bleo-exposed rats at day 10, while treatment with Dex or Cpd A repressed this increase. Inflammatory mediators were reported to increase in either bronchoalveolar lavage or homogenized lung tissues from BPD (3, 18, 19, 20). In this study, the mRNA expression of IL-1β and CCL2 significantly decreased in Dex-treated rats. Cpd A also led to a significant decrease in CXCL1 and CCL2 gene expression. The observed increase following Bleo exposure disappeared at day 14, and mRNA levels of inflammatory mediators decreased soon after exposure to Bleo.

Furthermore, CD68-positive cells were increased in Bleo-exposed lungs. This result is consistent with previous studies (21, 22, 23). However, the number of CD68-positive cells decreased in lungs treated with Cpd A. In this study, Ly6G-positive cells also increased in Bleo-exposed rats, and decreased in rats treated with Cpd A. These results suggest that lung inflammation is reduced upon treatment with Cpd A. GCs were previously reported to repress inflammation and to improve the severity of BPD (18). Cpd A is also worth investigating for this ability to attenuate the severity of BPD, in addition to GCs.

Here, we demonstrated that the level of TGF-β1 in the lungs of Bleo-exposed rats was increased on day 14 and that Cpd A treatment suppressed this increase. TGF-β plays a key role in tissue development and injury repair (24). However, overexpression of TGF-β in neonatal mice causes morphometrical changes in the lungs that are similar to those observed in BPD (25). In bronchoalveolar lavages from preterm babies with lung injury, the level of TGF-β was shown to increase and to correlate with the severity of BPD; however, inhibition of TGF-β activity improved alveologenesis in hyperoxia-exposed mice lungs (26). We saw an increase in TGF-β1 mRNA on day 14, and found that Cpd A treatment suppressed the increased expression of TGF-β1, similar to Dex-treatment. This suggests that Cpd A has the potential to relieve developmental arrest and to attenuate lung injury. In this study, the level of TGF-β1 in Bleo-exposed rat lungs was lower than controls on day 10. Martin and colleagues reported that the level of TGF-β in hypoxia-exposed mice was lower than in normoxic mice during the early stages (27). They suggested that the expression of TGF-β in injured lungs initially decreases and then increases during the chronic stage (27).

We found that the average body weight of rats in the Dex group on day 14 was significantly lower compared with the other groups. GCs are known to cause growth retardation by suppressing the growth hormone/insulin-like growth factor-1 axis (28). Furthermore, GCs induce muscle degradation (12) and loss of bone density (29). Consequently, Dex-treated pups gained less weight. Conversely, Cpd A has not been reported to have negative effects on bone and muscle (12, 29) and our results are consistent with these previous reports.

Recently, some SEGRMs, such as ADZ7594 and Mapracorat, have been approved for use in clinical trials. ADZ7594 is reported to improve lung function and symptoms in adult patients with asthma without any severe adverse effects (30). Mapracorat is used as a topical treatment for inflammatory skin disorders (31) and has been reported to reduce the severity of psoriasis without any serious adverse effects.

This study has several limitations. First, Dex and Cpd A treatment was initiated on the same day as the bleomycin exposure began. Thus, Dex and Cpd A were used as a prophylactic medicine in this study. In clinical practice, GCs are only administered to infants suffering from severe BPD. It is necessary to study whether Cpd A has efficacy during chronic stages of BPD. Second, in this study, we only evaluated body weight for adverse effects. Impaired neurodevelopmental is a significant adverse effect of GC treatment, particularly in children. Further investigations are needed to evaluate neurodevelopment following Cpd A treatment.

In conclusion, our results indicate that Cpd A decreases lung inflammation and improves lung morphometric changes in an experimental model of BPD. Although further study is needed, our results suggest that SEGRMs, including Cpd A, may be promising candidates for the therapy of BPD.

## ACKNOWLEDGEMENTS

The authors would like to thank Kazumi Honda and Yumiko Yoshimura for assistance with the biochemical analysis, and, Yusuke Onishi and Yasuhiro Ogino for assistance with the immunohistochemical analysis.

## FIGURE LEGENDS

Supplemental figure 1. GCs bind to the GRs in the cytoplasm. After ligand binding, the GRs translocate into the nucleus, where they regulate gene transcription with two processes: transactivation and transrepression. In transactivation, a GR-homodimer binds to glucocorticoid-response elements (GREs) and induces the expression of anti-inflammatory genes. In transrepression, the monomeric GRs interfere with other transcription factors associating with inflammation and inhibit the expression of pro-inflammatory gene. SEGRMs act as a ligand for GRs and have anti-inflammatory properties via only transrepression.

Supplemental figure 2. Study protocol. Bleomycin (Bleo) or saline was injected peritoneally from day 0 to day 10 of life. Vehicle (distilled water), Dexamethasone (Dex), or Compound A (Cpd A) were injected until day 13. 1) Control group; Normal saline-exposed and vehicle-treated, 2) Bleo group; Bleo-exposed (1mg/kg) and vehicle-treated, 3) Dex group; Bleo-exposed and Dex-treated (0.1mg/kg), and 4) Cpd A group; Bleo-exposed and Cpd A-treated (1mg/kg).

Pups were sacrificed for PCR at day 10, and for morphological studies and PCR at day 14.

