## Supplementary figures and images for "A selective glucocorticoid receptor modulator attenuates lung inflammation and improves alveolarization in a neonatal rat model of bronchopulmonary dysplasia"

### Supplemental Figure 1

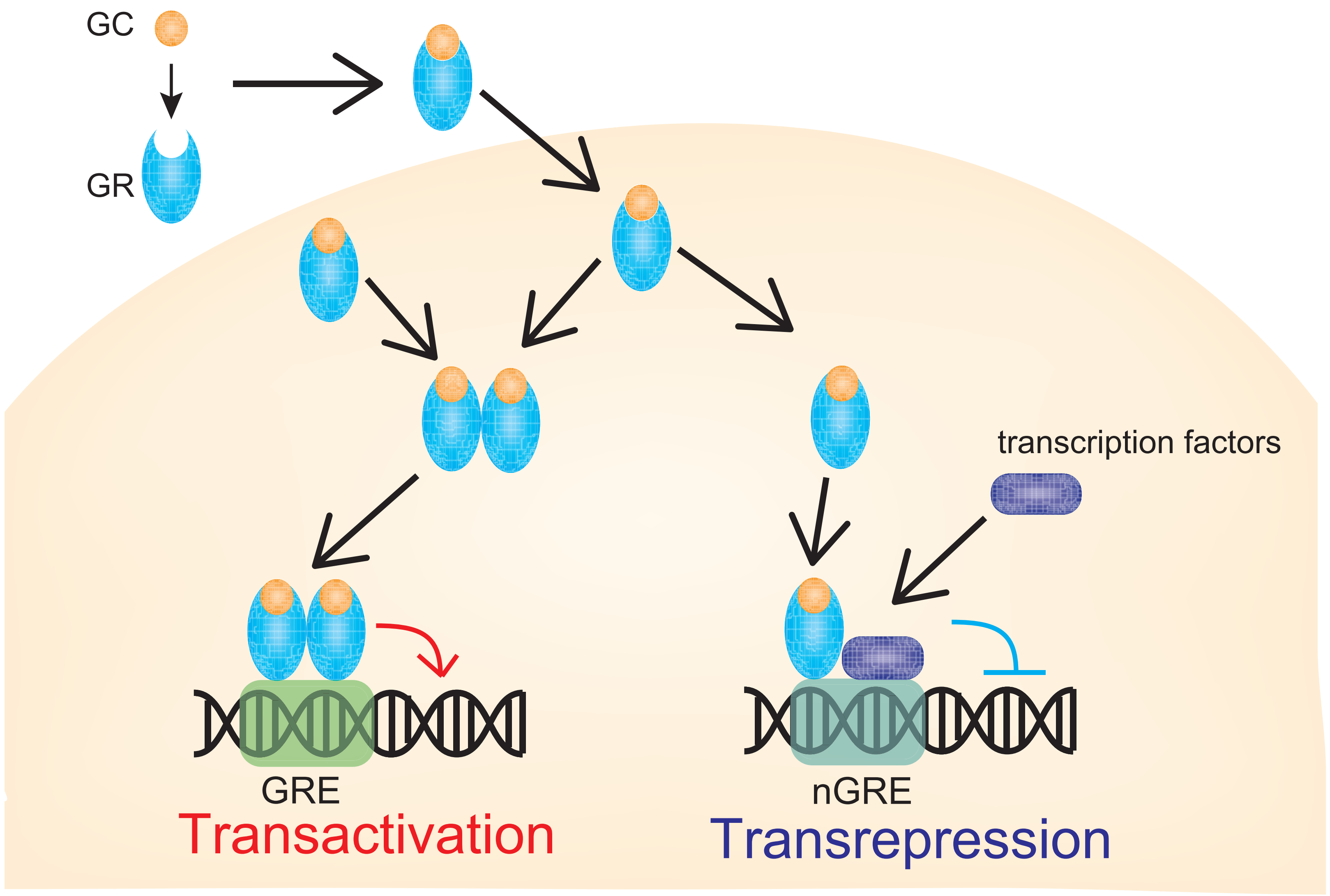

### Supplemental Figure 2

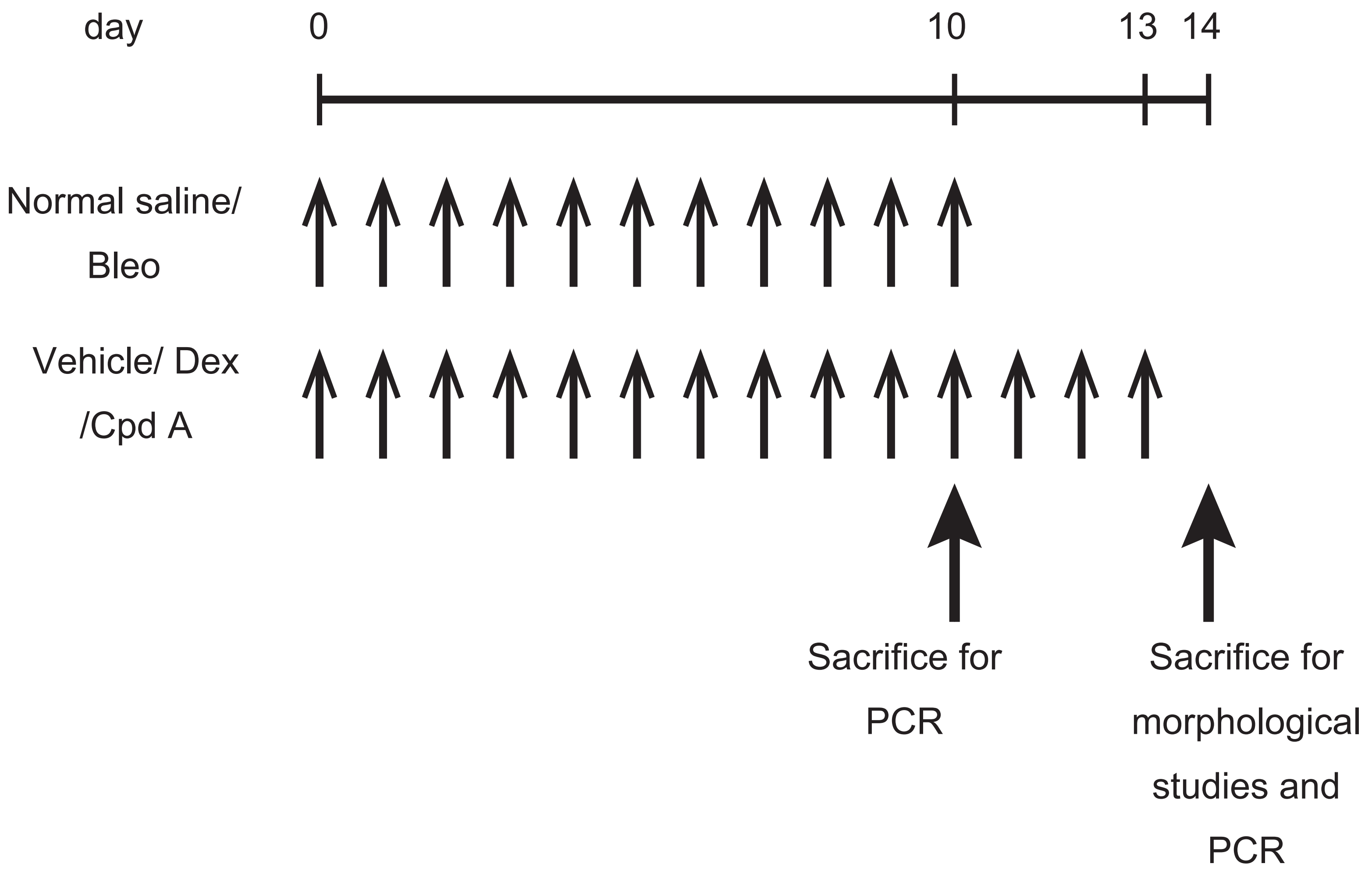
